# The Impact of Different Learning Processes on Acquisition, Transfer, and Proprioception in Complex Motor Tasks

**DOI:** 10.1101/2025.10.29.684950

**Authors:** Reshma Babu, Hannah J. Block

## Abstract

Motor skill learning involves multiple mechanisms, including use-dependent learning (UDL), reinforcement learning (RL), and error-based learning (EBL). These operate over different time scales and neural pathways, contributing uniquely to skill acquisition, consolidation, and transfer. Here we asked how these mechanisms support the acquisition of a spatially complex maze navigation task in five groups of healthy young adults. Groups received one of five types of feedback during training of their unseen dominant hand: UDL (no feedback), RL (binary success/failure feedback with a static threshold), RLA (binary feedback with an adaptive threshold), RLB (binary feedback with adaptive threshold and brief flash of cursor feedback), or EBL (continuous real-time cursor feedback). Skill, transfer, and proprioceptive acuity were assessed pre- and post- training using a speed-accuracy function (SAF) for each hand and a two-alternative forced-choice shape discrimination task. Results showed that UDL and RL groups exhibited no improvement post-training, while EBL, RLA, and RLB groups demonstrated accuracy improvements. EBL and RLB participants experienced a significant reduction in movement variability, with EBL showing a greater decrease compared to UDL. The left-hand SAF revealed improvements in accuracy across all groups except UDL. All groups showed reduced variability in the left hand, suggesting intermanual transfer, with EBL transferring more variability improvements than UDL. No significant proprioceptive changes were observed in any group. These findings provide new insights into motor skill learning, emphasizing that even minimal feedback can facilitate complex skill acquisition and transfer and has significant implications for studies where error-based learning may not be applicable.

**New and Noteworthy:** Minimal, adaptive binary feedback can effectively support the acquisition and transfer of spatially complex motor skills. While use-dependent and static reinforcement learning failed to enhance performance, adaptive reinforcement and error-based feedback significantly improved accuracy and reduced movement variability. Notably, these gains transferred to the untrained hand, highlighting the potential of reinforcement-based strategies in contexts where error-based learning is limited or unavailable, offering important implications for rehabilitation and motor training design.

## Introduction

Learning any motor skill likely involves multiple underlying processes. These include use-dependent learning, reinforcement learning, error-based learning, and strategy use (Spampinato & Celnik, 2021). Each of these processes depends on a distinct set of neural substrates with differing timescales of action that usually work together to account for skill learning. Depending on the task complexity and the stage of learning being considered, the relative contributions of each process may differ (Spampinato & Celnik, 2021). However, much of the literature focuses on relatively simple motor skills, limiting our understanding of how these processes can be used to learn more complex skills. Simple tasks may be characterized by consistent timing, minimal cognitive demand (Wulf & Lewthwaite, 2009), and fewer degrees of freedom, often resulting in predictable trajectories with short and steep learning curves (Schmidt & Lee, 2011). Complex tasks may require coordination across multiple joints and muscle groups (Kawashima et al., 2005; Swinnen & Wenderoth, 2004), leading to more variable movement patterns. These tasks engage closed-loop control, necessitating real-time feedback, higher cognitive processing (Wulf & Lewthwaite, 2009), and adaptive strategies for planning and error correction (Newell & Boucher, 1974; Schmidt & Lee, 2011).

Error-based learning (EBL) involves receiving online and offline feedback to modify behavior based on an error signal: the difference between the expected sensory outcomes of movement and the actual sensory outcomes. Providing error feedback while performing a task gives enough information as to why a past movement could have failed and what should be corrected so that the goal of the movement can be achieved (Tseng et al., 2007). These sensory prediction errors can be used to calibrate and update the internal representations of the body and the environment (Block & Bastian, 2011; Mazzoni & Krakauer, 2006). For example, when icing a cake, we refine our technique through a process of feedback and adaptation to achieve the best results. Initially, the appearance of the cake and icing sets our expectations. As we begin applying the icing, we observe how it interacts with the cake. If we notice uneven areas or crumbly spots, this feedback tells us where adjustments are needed. With each attempt, we refine our movements, reduce errors, and achieve a smoother, more polished finish.

Reinforcement learning (RL) occurs in response to binary feedback about the movement’s success or failure (Sutton & Barto, 1999). RL mechanisms are in play when we practice making a basket or trying to shoot a free kick where the outcome of importance is whether you make the shot or not (success or failure). The RL process involves preferentially selecting movements that maximize the chance of success. This requires exploration to determine the movements that are associated with success vs. failure. RL is driven by a prediction error that relates to the probability of a behavior resulting in success. This is essentially a credit assignment problem-different actions are assigned credit based on their success rate and the brain modifies behavior to increase the chance of success (A. S. Therrien et al., 2016a). Studies looking at RL in a simple motor task have found that having an adaptive threshold for success, making success gradually harder to achieve, increases the chances of the feedback being helpful and aiding the movement (Schultz, 1998). In a more complex task involving pattern matching for length and slant, when subjects had to match a straight trajectory that was slanted in depth, learning was observed when a composite feedback based on the slant and length error was provided (Kooij et al., 2021). However, feedback on one factor did not improve the other, and increasing the number of factors and complexity of the task reduced the chance of learning the task, as when subjects were asked to match a curved trajectory where the composite feedback was based on additional factors like curvature and smoothness of the trajectory (Kooij et al., 2021).

Use dependent learning (UDL) involves practicing the same movement repeatedly, not requiring any feedback of the actual movement or its outcome. Studies have shown that repetition reduces variability in movement (Huang et al., 2011; Verstynen & Sabes, 2011) and movement time (Hammerbeck et al., 2014) by improving movement planning (Mawase et al., 2018). UDL is thought to elicit the plasticity mechanisms in the motor cortex (M1) that bring about long-term learning effects. In real life motor skill learning, UDL would be involved when athletes practice the same throw, shot or free kick repeatedly or musicians play the same piece repeatedly.

Performance on multiple measures can reflect skill learning. Performance on a skill is limited by task difficulty, with the goal being to improve speed and accuracy (Dayan & Cohen, 2011; Krakauer & Mazzoni, 2011). A shift in the speed-accuracy tradeoff is a standard measure of skill gain (McGrath & Kantak, 2016; Reis et al., 2009; Shmuelof et al., 2012), reflecting that accuracy has improved without sacrificing speed or vice versa. Skill can also be acquired by reducing the intrinsic variability and noise (Shmuelof et al., 2012) by moving from exploring various strategies to sticking with the best strategy (Sternad et al., 2014). Skill is known to transfer from the trained arm to the untrained arm, though there are inconsistent results on the role of arm dominance in transfer (Halsband & Lange, 2006; Taylor & Heilman, 1980). This diversity and asymmetrical features of transfer are believed to be task dependent (Yadav & Mutha, 2020). Information can be transferred through visuospatial mechanisms, making use of extrinsic spatial coordinates or stimulus sequence, and through body-related effector transfer, taking advantage of homologous neuronal and muscular motor components of the untrained side.

Motor skill learning has also been linked to improvements in proprioception. Practicing a simple center-out reaching task showed improvements in precision of estimating the hand position (J. D. Wong et al., 2011). Adding binary feedback about the knowledge of results to the task improved proprioceptive acuity as well (Bernardi et al., 2015). A recent study by Mirdamadi and Block found both neurophysiological and behavioral somatosensory changes associated with learning a spatially complex maze-tracing skill (Mirdamadi & Block, 2020). Short-latency afferent inhibition (SAI), which is indicative of the projections from S1 to M1, was measured after training on a maze tracing skill. It was found that SAI increased with training, along with proprioceptive acuity (Mirdamadi & Block, 2020).

The present study asks whether a spatially complex skill like maze navigation can be acquired using use-dependent and reinforcement mechanisms and how the learning, intermanual transfer and associated proprioceptive changes would compare to error-based learning. Participants were randomly assigned to learn the skill by use-dependent mechanisms alone (UDL group), by reinforcement with a static or adaptive threshold for success (RL and RLA groups), by reinforcement with a brief moment of online error feedback (RLB group), or with full online error feedback (EBL group).

## Methods

### Participants

A total of 115 subjects between the ages of 18-45 participated in the study (72 male, mean age 23 ± 5.65). Subjects were randomly assigned to one of the five groups. All subjects were right-handed as per the Edinburgh handedness questionnaire (Oldfield, 1971) with normal or corrected-to-normal vision. They reported no musculoskeletal injuries, neurological diseases, or attention disorders. The protocols were approved by the Institutional Review Board (IRB) of Indiana University Bloomington and were performed in accordance with the IRB guidelines and regulations.

### Experimental Design

Each participant completed one session. The session took around 2 hours with every task taking 12-15 mins. Subjects were given scheduled 2-3 min breaks between every task to minimize fatigue. The session began with the pre-training assessment of skill, transfer, and proprioception, in a random order. These were followed by training on the maze navigation task, that differed based on random group assignment. Finally, the skill, transfer and proprioceptive assessments were repeated post-training in a random order.

All tasks were performed using the KINARM Endpoint 2D robotic manipulandum (BKIN) (Fig 1A.). Participants were instructed to grasp the manipulandum with their hand while performing each task. The maze and all visual feedback were displayed on a horizontal screen, viewed through a horizontal mirror making the images appear to be in the plane of the manipulandum, while preventing subjects from having direct vision of their hand or the robot. Subjects were instructed that the center of the manipulandum corresponds to their hand location while performing the tasks. Data about hand movements including timing data and end point locations were recorded by the KINARM system for analysis.

**Fig 1:**
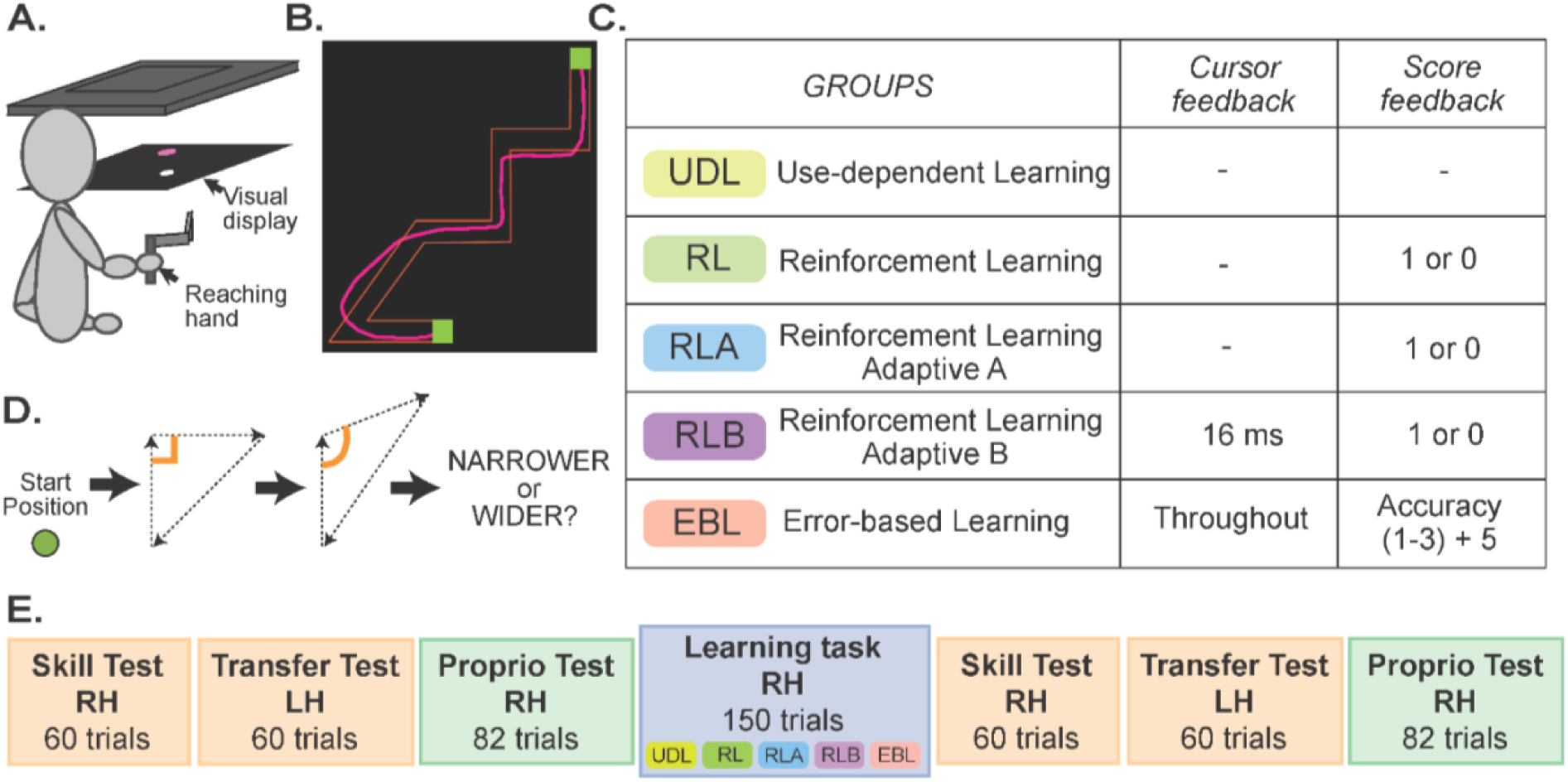
**A.** Experimental setup. 2D VR apparatus used for the motor and proprioceptive tasks. The task image was displayed on a horizontal television (top) and viewed in the mirror below. Not pictured: a drape over their shoulders that blocked vision of their hands. Subjects grasped the robotic manipulandum handle, below the mirror, throughout each task. Depending on the condition subjects saw a cursor indicating the location of their hand. **B.** Maze navigation task illustration. **C.** Feedback during skill learning consisted of cursor and/or points, depending on random group assignment. **D.** Display sequence for the shape discrimination task. Each trial began with the robot moving the hand to a start circle. The robot then moved the hand along the sides of an invisible triangle which was Target 1. Immediately after, a second triangle was formed (Target 2). After forming Target 2, the words NARROWER OR WIDER were displayed and subjects reported aloud to the experimenter whether the highlighted angle of Target 2 was narrower or wider than Target 1. **E.** The session began with the 3 pre-training assessments: skill, transfer, and proprioception, in random order. This was followed by training on the maze skill. Conditions of the skill learning task were determined by random group assignment. The session was concluded with the three post-training assessments: skill, transfer, and proprioception, in another random order.

### Skill learning task

For the motor skill, participants were asked to grasp the manipulandum handle with their right hand and navigate through an irregular shaped track (20×20 cm space, 1.5 cm width: Fig 1B) as accurately as possible within the desired movement time range (Kantak et al., 2018; McGrath & Kantak, 2016; Mirdamadi & Block, 2020). Each trial started with a red start-position square. 1s after moving the hand into the square, it turned green and the full track was displayed as a red outline with another green square at the end, signifying the start of the trial. Visibility of a cursor representing hand position was determined by group assignment. While moving at the right speed range, subjects were asked to be as accurate as possible in tracing the maze. After each movement, participants received feedback on their speed (too slow, too fast, good speed) and accuracy depending on their group. If given, accuracy feedback was provided only for the trials at the appropriate speed. During motor training, participants trained at a fixed movement time (MT) range (850-1100ms) for 150 trials. Only the trials in which subjects moved in the required speed range were included in the 150 trials. Each group received specific forms of feedback (Fig 1C.):

A. Use-dependent Learning (UDL): Subjects did not see their cursor after their start target turned green and did not receive any feedback on their accuracy. They only received speed feedback to ensure they moved in the appropriate speed range to complete the 150 trials.
B. Reinforcement Learning (RL): Subjects did not see their cursor throughout the trial but received binary feedback-1/0 based on their percentage accuracy. The threshold for performance accuracy remained at 35% throughout the experiment.
C. Reinforcement Learning Adapted (RLA): Subjects did not see their cursor throughout the trial, but they received binary feedback-1/0 based on percentage accuracy. The threshold for performance accuracy started at 35% and it increased by 5 when subjects moved accurately for 3 out of 5 trials and decreased by 3 when they were not accurate.
D. Reinforcement Learning B (RLB): Identical to RLA, but subjects also saw their cursor for 16 ms at the center of the maze.
E. Error-based learning (EBL): Subjects saw a cursor representing their hand position (white circle, 10 mm diameter) throughout the session and received points based on their in-track accuracy. The visual cursor specifically appeared at the center of the manipulandum handle, which was surrounded by the fingers and thumb in a whole-hand grasp. They received 5 points for completing the maze and then additional points based on accuracy-3 points if they were 90% accurate, 2 points if they were 80% accurate and 1 point if they were 70% accurate (Mirdamadi & Block, 2020).

The subjects were asked to end accurately in the second target square at the end of the maze to successfully complete a trial. The trial ended when subjects entered the end target for EBL or an invisible 20 cm rectangle target at the same location of the end square target for all the other groups.

### Skill Assessment

Participants had to navigate the same maze with visual cursor feedback for skill assessment. The motor skill assessment looked at performance over six movement time (MT) ranges in random order (Kantak et al., 2018; McGrath & Kantak, 2016; Mirdamadi & Block, 2020), with 10 trials in each range (MT1: 350-600 ms; MT2: 600-850 ms; MT3: 850-1100 ms; MT4: 1100-1350 ms; MT5: 1350-1600 ms; MT6: 1600-1850 ms). Subjects did not receive any accuracy score but only speed feedback to successfully complete trials.

### Transfer Assessment

The skill assessment task was done exactly as the skill assessment task but with the left hand to assess intermanual transfer of learning.

### Proprioceptive Assessment

Proprioception was assessed using a passive two-alternative forced choice task (Martin et al., 2013; Mirdamadi & Block, 2020). Subjects were told that the center of the robotic manipulandum handle, which was grasped in the right hand, corresponded to hand position, and that the robotic manipulandum would passively move their hand to create different triangles in the workspace.

For each trial, the robot presented two triangles consecutively and the subject was required to determine the difference between the first and second shapes (Fig. 1D). Participants had to report whether the first angle of Target 2 was wider or narrower than Target 1. The trial sequence began with the robot moving the subject’s hand to the center start location, and after a 0.5-s pause, moving the subject’s hand around the contours of a 10-cm triangle starting from the bottom vertex of the shape so that each side took 1000 ms to complete, forming Target 1. Immediately after the presentation of the template shape, the robot presented Target 2. The robot then paused, and the computer displayed Narrower or Wider, signaling the end of the movement after which subjects verbally indicated their choice which was recorded by an experimenter with a button press on a standard keyboard.

The angles assessed ranged from 60°-120° and the step size was 2.5. Every set involved a standard (90°) and comparison target-the order of which was randomized. Subjects completed a total of 82 trials. There were two staircases in play-one where the comparison target started at a 60° triangle and increases towards 90° and the other where the comparison target started at a 120° triangle and decreases towards 90°. Every correct choice reduced the gap between the template and test target by 2.5° and every wrong choice increased the gap by 2.5°. There were 4 different trial types possible depending on the order of the standard and comparison target and which staircase it belonged to, which were arranged pseudo randomly. Each staircase consisted of 40 trials where 20 trials had the standard appear first and 20 had the comparison target appear first and 2 trials in the beginning to ‘start’ the staircase leading to 84 trials per session.

### Data Analysis

Data analysis was performed with MATLAB 2022b. For the proprioceptive data, the fitting was performed using the psignifit toolbox version 4.0 for MATLAB (Wichmann & Hill, 2001). The data and analysis codes used to produce this manuscript are publicly available at https://osf.io/j6f7e/.

### Maze tasks

The movement time (MT) and cursor coordinates throughout the trial were recorded for each trial. Only trials in the correct MT range were analyzed. The in-track accuracy was calculated as the percentage of points inside the maze with respect to the total number of points in the trajectory made by the hand (Fig. 1B). Variability of the movement for each trial was also calculated as the root mean square distance of the trajectory from the bounds of the maze. For the learning task, performance was measured as a function of in-track accuracy and variability over time. 15 bins of 10 trials each were compared to check for practice effects on performance-medians computed both for accuracy and variability. For the assessment tasks, median speed-accuracy trade-off functions and the median variability of movement at each speed range were plotted pre and post learning. A composite score was computed by averaging accuracy or variability across the speed ranges.

### Proprioceptive task

For each assessment, the proportion of trials that participants responded “narrower” or “wider” for different test shapes was calculated. The data was fitted with a logistic function upon which the just noticeable difference (JND) and point of subjective equality (PSE) was calculated (Gescheider et al., 1985). JND, which represents the proprioceptive sensitivity, is half of the difference between the x-axis values yielding a 25% and a 75% response on the function. PSE representing bias was the x-axis value yielding a 50% response on the function. The PSE and JND before and after performing the learning task were compared.

### Statistical Analysis

All the statistical analysis was done using IBM SPSS Statistics (Version 29) and JASP (Version 0.19.1). The effect of practice on the accuracy and variability of performance on the maze was evaluated using a mixed-model ANOVA with Group (UDL, RL, RLA, RLB, EBL) and Block (Block 1, Block 15) being the factors. The training effects on proprioceptive bias and sensitivity were evaluated with a mixed model ANOVA with Group (UDL, RL, RLA, RLB, EBL) and the different time points (pre, post) as factors. Pre-training accuracy and variability was subtracted from the post-training values and then compared across the six different speed ranges using a mixed model ANOVA with Speed (1-6) and Groups (UDL, RL, RLA, RLB, EBL) as factors. Post-hoc analysis was done on the significant effects using Bonferroni correction. The changes were also compared to zero to identify whether there were significant differences at each speed using a one-sample t-test. The composite scores were compared across groups using a one-way ANOVA and post-hoc tests were done on significant differences. One-sample t-tests were also done to check if the composite score was different from zero for each group. Alpha was defined as 0.05 for all hypothesis tests.

## Results

### Motor acquisition

Motor acquisition can be quantified as the improvement in performance (accuracy and variability) over time while practicing the task. The example subject in Fig. 2A-B had an increase in accuracy and decrease in variability over time, consistent with progress in acquiring the skill.

**Fig 2:**
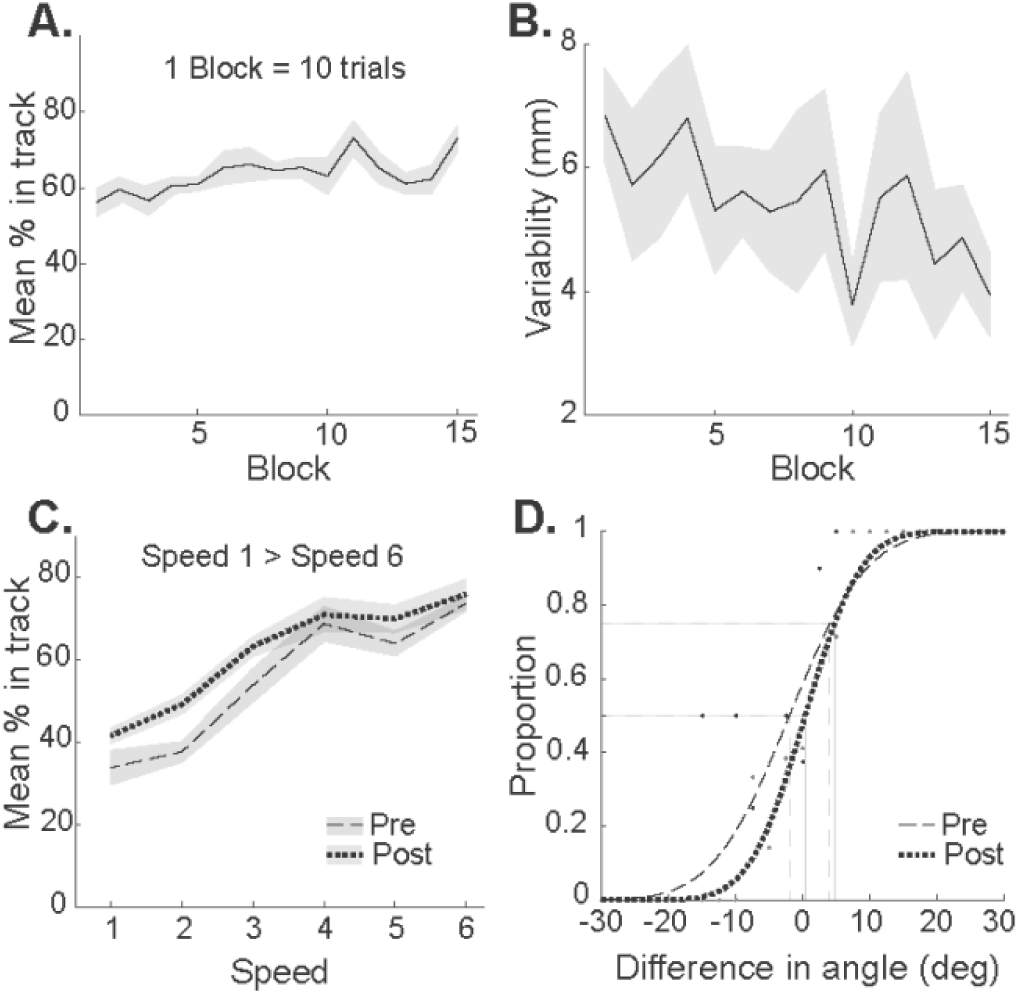
Example subject. **A:** Mean in-track accuracy for an example RLA subject over time with each block equaling 10 trials. **B:** Variability in hand movement calculated as the root mean square of the distance of the trajectory from the maze averaged for every 10 trials. **C:** Pre (dashed line) and post (dotted line) speed-accuracy trade-off functions where speed 1 is the fastest and speed 6 the slowest. This subject shows improvement in skill after learning (higher accuracy for a given speed). A similar graph can be made for the transfer assessment task as well. **D:** A sigmoid function that indicates the PSE which is the value of the stimulus axis (difference in angle between the comparison and standard stimulus) corresponding to proportion comparison of 0.5 and JND which equals the difference between the values on the stimulus axis corresponding to 0.75 and 0.5. This is an example subject whose PSE and JND gets better with time.

During the 15 blocks of training trials, we observed some learning (improvement in in-track accuracy) in the EBL group (59.95% to 64.40% accuracy), while the performance of RL (41.79 to 38.45%), RLB (47.78 to 41.92%), RLA (41.96 to 31.39%), and UDL (43.19 to 34.50%) groups became worse by this measure (Fig. 3A). Comparison of in-track accuracy in blocks 1 vs. 15 across groups (early and late in training) revealed a main effect of Block (F (1,110) = 21.514, p < 0.001, 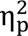 = 0.164) and Group (F (4,110) = 37.979, p < 0.001, 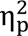 = 0.580) and an interaction effect (F (4,110) = 10.281, p < 0.001, 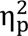 = 0.272). This indicates that the groups changed differently between the two timepoints. Post-hoc analysis of the main effect of group using Bonferroni correction showed that the EBL group had significantly better overall accuracy than every other group (all p < 0.001), which is to be expected since only EBL had continuous cursor feedback. In addition, the RLB group had significantly better performance than UDL (p = 0.015), and RLA (p < 0.001). The post-hoc analysis on the interaction effects showed that the performance accuracy significantly improved across blocks for EBL (p = 0.002) while it significantly reduced for UDL (p < 0.001), RL (p = 0.047), RLA (p < 0.001) and RLB (p = 0.046).

**Fig 3:**
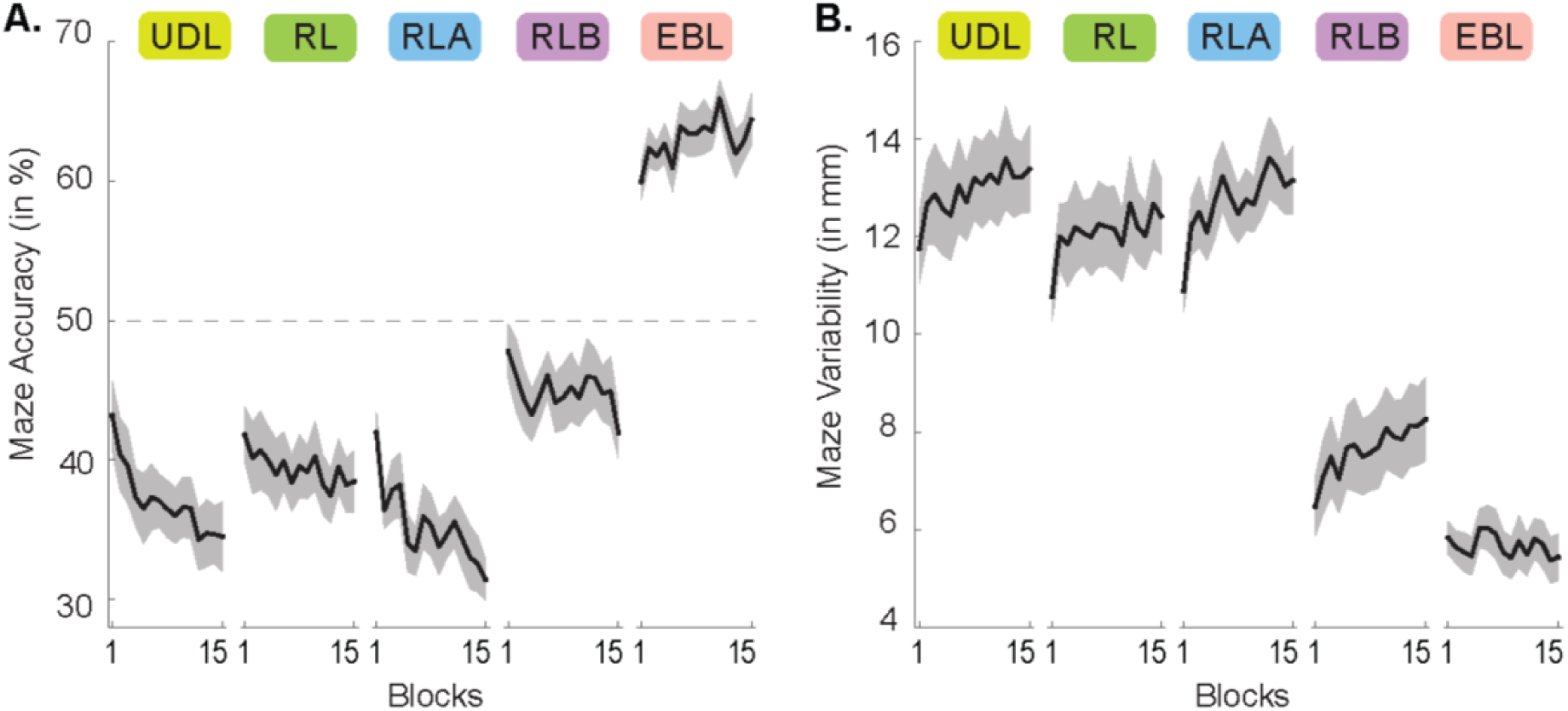
Performance during training. Comparison of mean in-track accuracy and variability for the five groups. **A:** For each group, mean percent in-track accuracy is plotted for the 15 blocks of training trials (black lines) with standard error of the mean (shading). **B:** For each group, mean movement variability (root mean square distance of the trajectory from the maze) is plotted for the 15 blocks of training trials with standard errors.

Movement variability (Fig. 3B) was low throughout training for the EBL group (5.83 mm to 5.43 mm). The other groups showed a slight increase in variability throughout training: RL (10.77 mm to 12.48 mm), RLA (10.89 mm to 13.16 mm), UDL (11.76 mm to 13.40 mm) and RLB (6.47 mm to 8.26 mm). Analysis of movement variability across timepoint (Block 1 vs. Block 15) and groups showed a main effect of Block (F (1,110) = 29.692, p < 0.001, 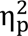 = 0.213) and Group (F (4,110) = 26.680, p < 0.001, 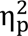 = 0.492) and an interaction effect (F (4,110) = 3.337, p = 0.013, η_p_^2^ = 0.108), suggesting the groups changed differently across time. The post-hoc analysis of the main effect of group (Bonferroni corrected) showed that the EBL group had lower movement variability than every other group (all p < 0.001), except RLB (p = 0.480) and the RLB group had lower movement variability than the UDL, RL, and RLA groups (all p < 0.001). The post-hoc analysis of the interaction effect showed that variability significantly increased across blocks as consequence of training for UDL (p = 0.009), RL (p = 0.013), RLA (p < 0.001), and RLB (p = 0.002) but did not significantly differ for EBL (p = 0.570).

### Skill learning

To assess gains in skill, the speed-accuracy and speed-variability functions were assessed for the right hand before and after training. The example subject shows an upward shift in the function post training which is consistent with learning (Fig 2C). Groups EBL, RLA, and RLB showed consistent evidence of gains in skill in terms of movement accuracy. Composite scores for accuracy in the right hand improved significantly after training in groups RLA (t (22) = 3.319, p = 0.003, Cohen’s d = 0.692), EBL (t (22) = 2.872, p= 0.009, Cohen’s d = 0.599) and RLB (t (22) = 2.698, p = 0.013, Cohen’s d = 0.563), but not in the RL group (t (22) = 0.529, p = 0.602, Cohen’s d = 0.110) or the UDL group (t (22) = −0.008, p = 0.993, Cohen’s d = −0.002) (Fig 4A bars). Considering the 6 speed ranges yielded further information about skill gain (Fig 4A lines, and Fig. S1). A mixed model ANOVA on the change in accuracy, with Speed (1-6) x Group (UDL, RL, RLA, RLB, EBL) as factors showed a main effect of group (F(4, 110) = 2.615 c, p = 0.039, η_p_^2^ = 0.87) and an interaction effect (F(16.159, 444.361) = 1.701, p = 0.043, η_p_^2^ = 0.58) but no main effect of speed (F(4.04, 444.361) = 2.086, p = 0.081, η_p_^2^ = 0.019). The sphericity assumption was verified using the Mauchly’s test, which was significant for speed, χ^2^ (14) = 66.625, p < 0.001. Therefore, the degrees of freedom were corrected using the Greenhouse-Geisser method, ε = 0.808. The post-hoc test on groups showed no difference.

**Fig 4:**
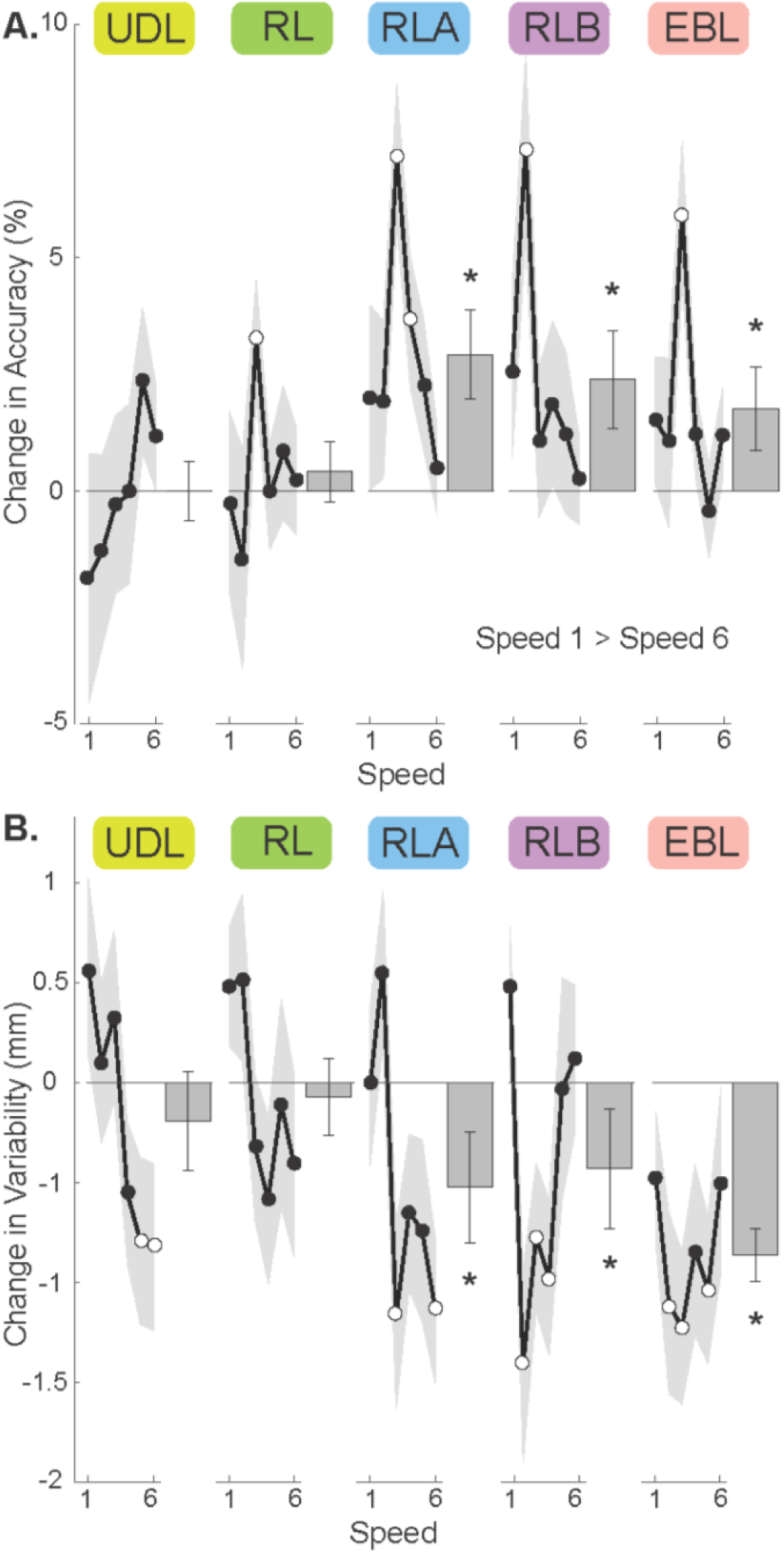
Skill gain. Change in the speedaccuracy (A) and speed-variability (B) trade-off functions. The six speeds are ordered from fastest (1) to slowest (6). Composite scores (averaged across speeds) are represented as bars. Open circles represent a change significantly different from zero. **A.** Positive values, reflecting increase in accuracy, indicate skill learning. **B.** Negative values, reflecting decrease in movement variability, indicate skill learning.

Comparing change in accuracy at each speed to zero revealed learning in at least one speed range for every group except UDL. Change in accuracy at Speed 3 was significantly different from 0 for RL (t (22) = 2.638, p = 0.015, Cohen’s d = 0.550), RLA (t (22) = 4.784, p < 0.001, Cohen’s d = 0.997) and EBL (t (22) = 3.872, p < 0.001, Cohen’s d = 0.807). Change in accuracy at Speed 4 was significantly different from 0 for RLA (t (22) = 2.562, p = 0.018, Cohen’s d = 0.534), and change in accuracy at Speed 2 was significantly different from zero for RLB (t (22) = 3.706, p = 0.001, Cohen’s d = 0.773). To understand the effect of group on change in accuracy at Speed 3 (the practiced speed), a one-way ANOVA showed a main effect of group (F (4,110) = 4.045, p = 0.004). Post-hoc tests suggest that UDL differed from RLA (p = 0.009) and EBL (p =0.046).

Groups EBL, RLA, and RLB also showed consistent evidence of skill gain in terms of movement variability. Composite scores for variability showed significant improvements for groups RLA (t (22) = −2.183, p = 0.040, Cohen’s d = −0.455), EBL (t (22) = −3.747, p < 0.001, Cohen’s d = −0.781) and RLB (t (22) = −2.440, p = 0.023, Cohen’s d = −0.509), but not RL (t (22) = −0.362, p = 0.721, Cohen’s d = −0.77) or UDL (t (22) = −0.936, p = 0.359, Cohen’s d = −0.191) (Fig 4B bar). A mixed model ANOVA on the change in variability with Speed (1-6) x Group (UDL, RL, RLA, EBL, RLB) as factors showed a main effect of speed (F (4.555, 501.033) = 3.912, p = 0.002, 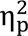 = 0.034) and an interaction effect (F (18.219, 501.033) = 1.848, p = 0.018, 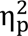 = 0.063) but no main effect of group (F (4, 110) = 2.062, p = 0.091, 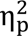 = 0.070). This indicates that the change in variability differed for the various speeds differently based on the groups (interaction effect) and also in general. The sphericity assumption was verified using the Mauchly’s test, which was significant for speed, χ^2^ (14) = 24.324, p = 0.042. Therefore, the degrees of freedom were corrected using the Greenhouse-Geisser method, ε = 0.911.

All groups showed variability improvements in at least one speed range, except for group RL. Change in variability at Speed 3 was significantly different from zero for RLA (t (22) = −2.540, p = 0.019, Cohen’s d −0.530), EBL (t (22) = −3.208, p = 0.004, Cohen’s d −0.669), and RLB (t (22) = −2.087, p = 0.049, Cohen’s d −0.435). Change in variability at Speed 2 was significantly different from zero for EBL (t (22) = −2.456, p = 0.022, Cohen’s d = −0.512) and RLB (t (22) = −2.917, p = 0.008, Cohen’s d = −0.608). Change in variability at Speed 5 was significantly different from zero for UDL (t (22) = 2.176, p = 0.041) and EBL (t (22) = −2.822, p = 0.010, Cohen’s d = −0.588).

Change in variability at Speed 6 was significantly different from zero for UDL (t (22) = −2.082, p = 0.049, Cohen’s d = −0.425), and RLA (t (22) = −3.112, p = 0.005, Cohen’s d = −0.649). Change in variability at Speed 4 was significantly different from zero for RLB (t (22) = −2.499, p = 0.020, Cohen’s d = −0.521). To understand the effect of group on change in variability at Speed 3 (the practiced speed), a one-way ANOVA showed a main effect of group (F (4,110) = 2.604, p = 0.040, Cohen’s d = 0.086). Post-hoc tests did not show any difference between specific groups.

### Transfer of skill

To assess intermanual transfer of skill, the speed-accuracy and speed-variability functions were examined for the untrained left hand before and after right-hand training.

Groups RL, RLA, and RLB and EBL showed consistent evidence of intermanual transfer in terms of movement accuracy. Change in composite accuracy scores for the left hand showed no group effect (F (4,110) = 1.243, p = 0.297), but the groups RL (t (22) = 2.347, p = 0.028, Cohen’s d = 0.489), RLA (t (22) = 3.396, p = 0.003, Cohen’s d = 0.708), EBL (t (22) = 4.439, p < 0.001, Cohen’s d = 0.926) and RLB (t (22) = 2.219, p = 0.037, Cohen’s d = 0.463) had significantly more improvement than zero (Fig. 5A bars). A mixed model ANOVA on change in accuracy for the left hand, with Speed (1-6) x Group as factors, showed a main effect of speed (F (3.982, 438.031) = 2.828, p = 0.025, 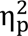 = 0.025). There were no group (F (4, 110) = 1.243, p = 0.297, 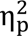 = 0.043) or interaction effects (F (15.928, 438.031) = 1.188, p = 0.274, 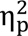 = 0.041) (Fig. 5A lines and Fig. S2). The sphericity assumption was verified using the Mauchly’s test, which was significant for speed, χ^2^ (14) = 62.768, p < 0.001. Therefore, the degrees of freedom were corrected using the Greenhouse-Geisser method, ε = 0.796.

**Fig 5:**
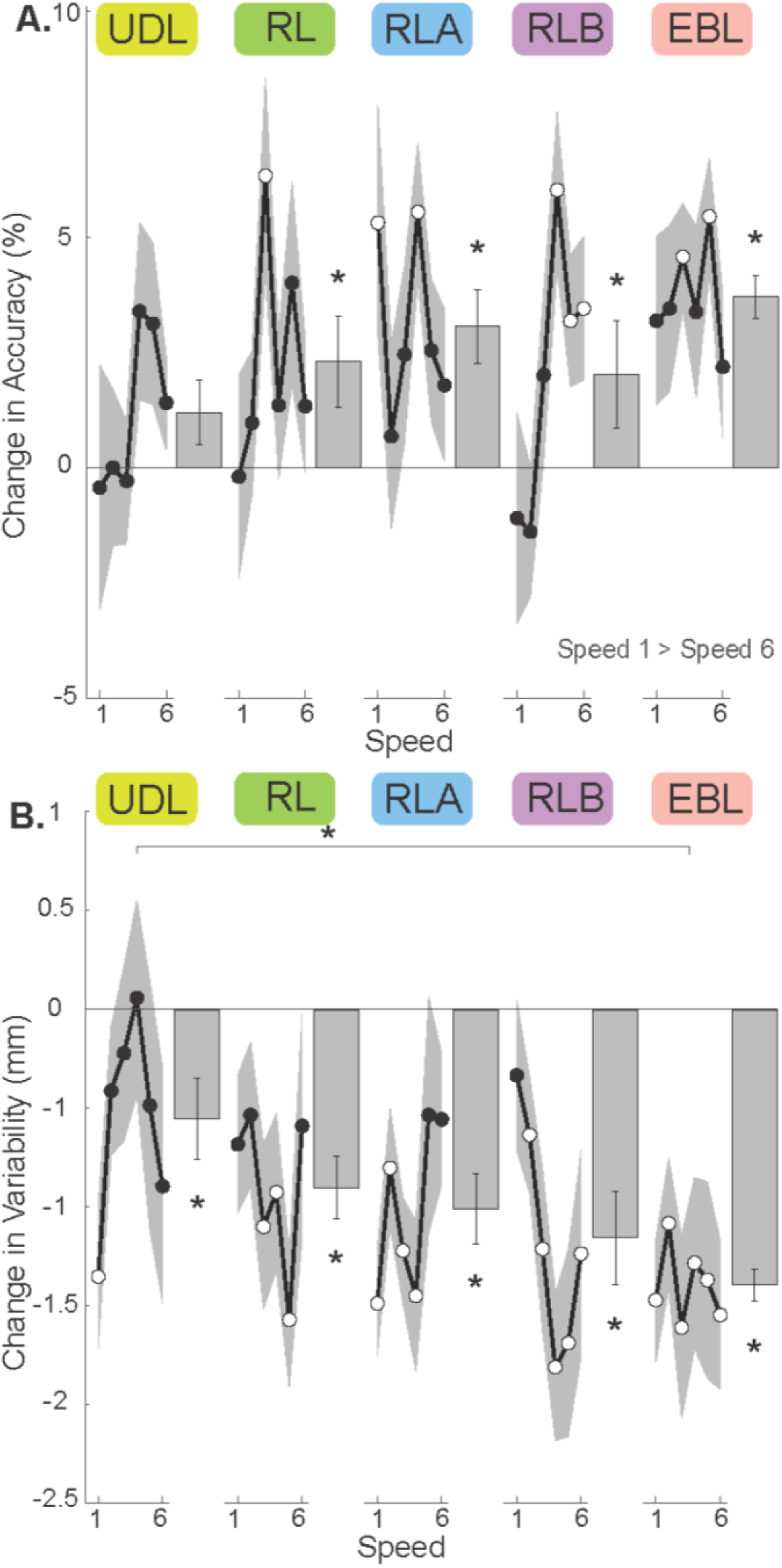
Intermanual transfer. Difference in speed-accuracy (A) and speed-variability (B) trade-off functions in the untrained left hand. The six speeds are ordered from fastest (1) to slowest (6). Composite scores (averaged across speeds) are represented as bars. Open circles represent a change significantly different from zero. **A.** Positive values, reflecting increase in accuracy, indicate skill learning. **B.** Negative values, reflecting decrease in movement variability, indicate skill learning.

All groups except UDL showed intermanual transfer of accuracy in at least one speed range. Change in accuracy at Speed 3 was significantly different from 0 for RL (t (22) = 2.978, p = 0.007, Cohen’s d = 0.621) and EBL (t (22) = 3.962, p < 0.001, Cohen’s d = 0.826). Change in accuracy at Speed 4 was significantly different from 0 for RLA (t (22) = 3.720, p = 0.001, Cohen’s d = 0.776) and RLB (t (22) = 3.569, p = 0.002, Cohen’s d = 0.744). Change in accuracy at Speed 6 was significantly different from zero for RLB (t (22) = 2.220, p = 0.037, Cohen’s d = 0.463), and Speed 5 for EBL (t (22) = 4.314, p < 0.001, Cohen’s d = 0.900) and RLB (t (22) = 2.231, p = 0.036, Cohen’s d = 0.465). Change in accuracy at Speed 1 was significantly different from zero for RLA (t (22) = 2.139, p = 0.044, Cohen’s d = 0.446).

All five groups showed significant intermanual transfer of learning in terms of movement variability. Change in composite variability scores for the left hand (Fig. 5B bars) showed a group difference (F (4,110) = 2.748, p = 0.032), with UDL and EBL groups differing from each other (p = 0.017). Change in composite variability score for the left hand was significantly different from zero in all five groups: UDL (t (22) = −2.184, p = 0.040, Cohen’s d = −0.455), RL (t (22) = −5.184, p < 0.001, Cohen’s d = −1.081), RLA (t (22) = −7.176, p < 0.001, Cohen’s d = −1.496), EBL (t (22) = −7.618, p < 0.001, Cohen’s d = −1.588) and RLB (t (22) = −6.627, p < 0.001, Cohen’s d = −1.382), consistent with intermanual transfer of skill. A mixed model ANOVA on change in variability with Speed (1-6) x Group as factors (Fig. 5B lines) showed a main effect of group (F (4, 110) = 2.748, p = 0.032, 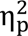 = 0.091). There were no speed (F (4.414, 485.547) = 0.756, p = 0.567, 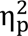 = 0.007) or interaction effects (F (17.656, 485.547) = 1.232, p = 0.232, 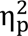 = 0.043). The post-hoc test showed that UDL differed from EBL (p = 0.020). The sphericity assumption was verified using the Mauchly’s test, which was significant for speed, χ^2^ (14) = 39.840, p < 0.001. Therefore, the degrees of freedom were corrected using the Greenhouse-Geisser method, ε = 0.883.

All five groups had significant improvement in left-hand movement variability in at least one speed range. Change in variability at Speed 3 was significantly different from 0 for RL (t (22) = −2.644, p = 0.015, Cohen’s d = −0.551), RLA (t (22) = −4.573, p < 0.001, Cohen’s d = −0.954), EBL (t (22) = −3.490, p = 0.002, Cohen’s d = −0.728) and RLB (t (22) = −2.964, p = 0.007, Cohen’s d = −0.618). Change in variability at Speed 2 was significantly different from zero for RLA (t (22) = −2.679, p = 0.014, Cohen’s d = −0.559), EBL (t (22) = −3.272, p = 0.003, Cohen’s d = −0.682), and RLB (t (22) = −2.161, p = 0.042, Cohen’s d = −0.451). Change in variability at Speed 1 was significantly different for UDL (t (22) = −3.706, p = 0.001, Cohen’s d = −0.773), RLA (t (22) = −5.548, p < 0.001, Cohen’s d = −1.157), and EBL (t (22) = −4.770, p < 0.001, Cohen’s d = −0.995). Change in variability at Speed 4 was significantly different for RL (t (22) = −2.327, p = 0.030, Cohen’s d = −0.485), RLA (t (22) = −3.813, p < 0.001, Cohen’s d = −0.795), EBL (t (22) = −2.970, p = 0.007, Cohen’s d = −0.619), and RLB (t (22) = −4.876, p < 0.001, Cohen’s d = −1.017). Change in variability at Speed 5 was significantly different from zero for RL (t (22) = −4.609, p < 0.001, Cohen’s d = −0.961), EBL (t (22) = −2.750, p = 0.012, Cohen’s d = −0.573) and RLB (t (22) = −3.570, p = 0.002, Cohen’s d = −0.744), and change in variability at Speed 6 was significantly different from zero for RLA (t (22) = 2.424, p = 0.024), EBL (t (22) = −4.107, p < 0.001, Cohen’s d = −0.856) and RLB (t (22) = −2.377, p = 0.027, Cohen’s d = −0.496).

### Changes in proprioception

The PSE and JND values obtained from the sigmoid function can be compared pre and post learning to understanding the effect of different learning types on the proprioceptive properties. The example subject in Fig. 2D showed improvement in both proprioceptive bias (PSE) and sensitivity (JND) for the shape discrimination task after skill training. A repeated measures ANOVA to test for group differences pre to post for PSE showed no significant effect of group (F (4,110) = 0.752, p = 0.559) but a significant effect of time (F (1,110) = 4.944, p = 0.028) and no interaction effect (F (4,110) = 0.633, p = 0.640). There was no significant effect of group (F (4,110) = 0.237, p = 0.917), time (F (1,110) = 3.179, p = 0.077), or interaction (F (4,110) = 1.801, p = 0.134) for the JND. These results are not consistent with any improvement in proprioception, and no group differences were evident (Fig. 6).

**Fig 6:**
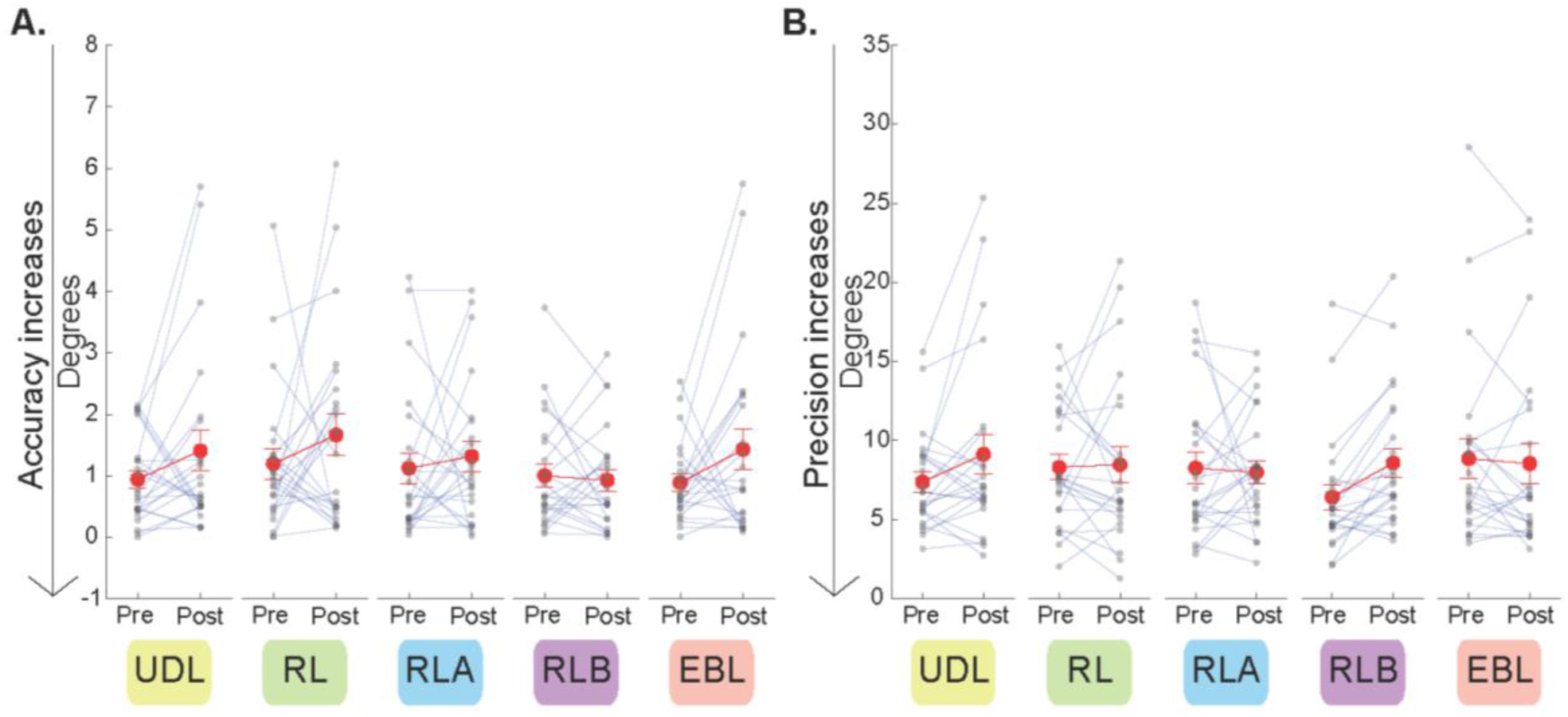
Proprioceptive bias (PSE) and sensitivity (JND) after motor skill learning (UDL, RL, RLA, RLB, EBL) **A.** PSE (point where 50% performance can be interpolated) pre- and post-training. A decrease in PSE post learning indicates an improvement in bias. None of the groups showed any evident improvement in bias. **B.** JND (smallest angle that can be reliably distinguished) pre- and post-training. A decrease in JND post learning indicates an improvement in sensitivity. No significant changes were evident.

## Discussion

Here we compared the effect of use-dependent, reinforcement, and error-based learning processes on a spatially complex motor skill task, evaluating skill acquisition, intermanual transfer of learning, and proprioception. Results suggest that skill learning, and transfer occurred robustly via error-based mechanisms (EBL). The reinforcement results were more mixed, but skill gains were evident for the RLA and RLB groups and intermanual transfer was evident for all three reinforcement groups. Results in the UDL group suggest that use-dependent processes alone may be insufficient to learn a spatially complex skill like maze tracing, but intermanual transfer can occur regardless.

### Performance during training depended on the learning type

Performing a complex task like maze navigation might require more than binary performance feedback about the success of the movement to show an immediate improvement in performance. There was a significant difference in the accuracy of subjects in the RLB group compared to UDL, RL and RLA, indicating that getting some cursor feedback along with the points feedback helped to perform such a maze navigation task much better than just binary feedback or in the absence of feedback. The subjects in RL and RLA groups also receive binary feedback but their accuracy in the maze navigation is comparable to the subjects in the UDL group meaning that the presence of the cursor even briefly may trigger error-based mechanisms that can supplement the reinforcement feedback presented, resulting in better performance.

The performance of subjects in RLB shows that binary feedback can be useful to learn a complex task like maze navigation when there’s a bit of additional information to supplement this binary feedback is provided. This is in agreement with previous studies that have found that binary feedback about a specific dimension can help reduce errors along that dimension (Kooij et al., 2021). However, continuous feedback throughout the trial may be most beneficial for the acquisition of more complex tasks: The accuracy in the EBL group where subjects saw their cursor throughout the trial was consistently greater than the RLB group where subjects briefly saw their cursor for 16 ms.

The variability in maze accuracy increased for the reinforcement groups as the trials progressed from Block 1 to Block 15. This could reflect more exploration of the task space, which is associated with reinforcement learning (Pekny et al., 2015; Sidarta et al., 2018, 2021) This increase in movement variability happened in all the groups receiving binary feedback irrespective of the type of threshold and whether they received the brief visual stimulus. The oscillatory behavior observed in the accuracy of UDL, RL, RLA are also indicative of a similar switching in strategy where subjects try various movements to try to succeed as it they try to explore the task space to identify what makes a movement successful and what does not (Sidarta et al., 2018).

It should be noted that the maze used for the assessment tasks was the same as the maze used during training with the varying learning types. All groups had continuous cursor feedback during the speed-accuracy function assessments, so subjects could have applied information they learned about navigating the maze during the pre-training assessment to the training task itself. However, this would have been a greater concern if all the groups had a similar performance during training; the poor accuracy of the no-feedback groups during training compared to the feedback groups is inconsistent with extensive carry-over effects from the assessment to the training.

### Skill gain was evident via reinforcement and error mechanisms

The RLA, EBL and RLB groups all showed robust skill learning, both in the speed-accuracy and the speed-variability relationships. There have been contrasting opinions on whether providing just binary feedback can aid learning a complex skill task. Kooij and colleagues found that for a pattern matching task, as the error can occur in many dimensions, singular feedback might not be enough to improve performance significantly across all dimensions (Kooij et al., 2021). But a recent study on locomotion has shown that a novel gait pattern can be learned and retained over 24 hrs. with reinforcement learning (Wood et al., 2024). This study found that despite increased variability which potentially affects upright stability, subjects learned a novel gait pattern using binary feedback about step length which they retained over 24 hrs.

In the current study, subjects receiving binary feedback based on an adapted criterion with or without the 16 ms feedback (RLB and RLA) showed improvements in skill that were similar to the group receiving both cursor and points feedback (EBL). In contrast, the RL group improved accuracy at only a single speed range and did not significantly improve movement variability. Together, these results suggest the adapted criterion for threshold in the RLA and RLB groups was an important driver of reinforcement mechanisms in this spatially complex motor skill task. This is consistent with literature involving simple skills usually reaching tasks or modified reaching tasks that have found that gradually increasing the threshold criterion improves the amount of learning and performance on tasks (A. Therrien et al., 2015; A. S. Therrien et al., 2016b).

While learning generalizes across speeds in a speed-accuracy function, studies have found that the speed that is practiced shows the greatest learning effect (Donchin et al., 2003; Shadmehr, 2004; Shadmehr et al., 2010). In the present study, skill gains were most prominent at Speed 3 for most of the groups which can be attributed to the 150 trials that subjects practiced at that speed. The learning effect was less at speeds that are further removed from the practiced speed, analogous to the generalization observed while practicing reaches to a target-some learning effect remains to the targets that are close to the practiced target, and this reduces as the spatial distance increases (Krakauer & Shadmehr, 2006; Shadmehr et al., 2010; A. L. Wong et al., 2016). The RLB group did not show significant improvement in accuracy at Speed 3 though they had a significant reduction in variability.

Results suggest that repeated practice alone is not enough to learn a complex skill task like maze navigation; performance improvements in the UDL group were limited to a single significant speed range at which movement variability improved for the trained hand. This control-policy might be too complex to learn exclusively by repeating a movement. This could be because although participants may use exploration strategies to explore the solution space, not receiving any feedback can lead to over exploration without any exploitation or even wrong movement patterns being reinforced (Sidarta et al., 2021). Repeated execution of the same movement may improve simple movements that do not require multi-joint coordination and multi-step movements well but does not work for complex skills. As task complexity increases, sequencing of movements, multi-joint coordination increases that would require more frequent and enhanced feedback (Boutin et al., 2012; Jarus & Gutman, 2001; Wulf et al., 1998).

### Intermanual transfer occurred by use-dependent, reinforcement, and error-based mechanisms

In the untrained left hand, the EBL and all three reinforcement groups improved on the composite accuracy score, suggesting robust intermanual transfer. All five groups improved on the composite variability score in the left hand, which indicates that even use-dependent processes alone could mediate transfer. For a complex skill task such as maze navigation, practice without feedback is enough to learn a basic idea of the movement that is required to successfully complete the task and to understand the underlying joint-pattern and speed-accuracy requirements and this transfers well to the other limb. This is also consistent with other studies on simple reaching that have found this right-left transfer (Stöckel et al., 2016; Yadav & Mutha, 2020). The addition of both points and cursor feedback improves the amount of transfer as this feedback aids to understand the solution space better and contextualize the movement to transfer better.

There can also be an effect of the coordinate system on the amount of transfer. Numerous studies have proposed that every skill can be represented as distinct and independent coordinate systeme (Criscimagna-Hemminger et al., 2003; Hikosaka et al., 1999; Lange et al., 2004; Shadmehr & Mussa-Ivaldi, 1994). The visually acquired information about the movement and targets are coded in an extrinsic, world-based reference frame while the muscular activation patterns are encoded in an intrinsic, body centered reference frame (Colby & Goldberg, 1999; Soechting & Flanders, 1989). The intrinsic representation using joints, musculoskeletal forces and dynamics (Criscimagna-Hemminger et al., 2003; Krakauer et al., 1999) accounts for the orientation of body segments relative to each other (Lange et al., 2004; Soechting & Flanders, 1989). Such a system is supposed to result in an effector-dependent representation of the skill that is not prone to transfer. Conversely, the extrinsic system refers the Cartesian coordinates of the task space, and they are an effector-independent representation which means they can improve in intermanual transfer (Hikosaka et al., 1999). However, there has been some recent studies regarding neural evidence for effector independent representation in intrinsic coordinates (Wiestler et al., 2014). The current paradigm tested transfer in the extrinsic space as the trajectory the subjects had to make remained the same across hands, which is probably why we see transfer across the different learning types. How transfer based on an intrinsic coordinate system (with a mirrored maze) would work is not completely known, but the effects would not be as prominent as in the current study. In many cases, the improvements in the left hand surpassed the magnitude of improvements in the trained right hand. This is likely due to the relatively worse initial left-hand performance. As all the subjects were right-handed, the left-hand had more scope for improvement (Shadmehr et al., 2015). The right-hand performance could have started out closer to asymptote, compared to the left hand, which has been observed in earlier studies (Krakauer, 2015; Wong et al, 2017).

### No changes in proprioception observed

Previous studies looking at proprioceptive differences following motor learning have used simple position discrimination at the center of the workspace either in the horizontal (J. D. Wong et al., 2011) or both horizontal and sagittal dimensions (Mirdamadi & Block, 2020). Since the present task involved a complex maze navigation through multiple corners of the maze, perception of angle might be particularly relevant. However, an angle discrimination task did not detect any effect of skill learning, nor any group differences in performance. This could be due to the greater complexity of the assessment as well as other methodological differences. Previous studies (Mirdamadi & Block, 2020; J. D. Wong et al., 2011) found a range of improvements in proprioceptive sensitivity (∼11% and 35%); Wong and colleagues used a task that involved comparing the location of their hand to a remembered position, relying heavily on working memory, while Mirdamadi & Block assessed proprioception with respect to a visual reference marker presented simultaneously. In the current experiment, subjects had to infer whether their test angle was wider or narrower than a target that appeared earlier, making the task both highly dependent on working memory and also more spatially and cognitively complex than the position assessments (Mirdamadi & Block, 2020; J. D. Wong et al., 2011). The proprioceptive angle perception task, though relevant to maze navigation, might have been too difficult as evidenced by the variability in individual performance. A proprioceptive assessment of lower difficulty not involving working memory might be needed to see clear proprioceptive differences associated with skill learning mechanisms.

### Implications and future studies

The effect of reinforcement on the acquisition of a skill seems to be particularly prominent in situations where sensory feedback is uncertain (Bernardi et al., 2015; Cashaback et al., 2017; Izawa & Shadmehr, 2011; Nikooyan & Ahmed, 2015), implying that this approach could be promising for rehabilitation, as patients with motor impairments often also exhibit sensory deficits (Connell et al., 2008; Hepworth et al., 2016). Consistently, reinforcement has been known to show improvements in motor learning in different populations of patients including cerebellar (A. Therrien et al., 2018; Vassiliadis et al., 2019) and stroke patients (Goodman et al., 2014; Quattrocchi et al., 2017; Widmer et al., 2022). In this study, it is clear that reinforcement can also be beneficial in spatially complex skill tasks involving multi-joint sequential movements.

UDL showing significant transfer (reduction in variability) means they can benefit from practice as well. This is consistent with existing research that focuses on repeated practice to improve movement after stroke and other movement disorders (Wang et al., 2020).

These results can be extended to other studies that involve a sequencing of movement elements to achieve the goal. It will also be interesting to further probe how other complex tasks involving varying cognitive or task demands, or timing information are affected by the various learning types. Future research on learning and transfer of spatially complex skills should investigate tasks involving multi-joint movements that can be broken down biomechanically to figure out whether elements like initial limb configuration and explicit awareness of posture can affect how skills are learned and transferred. Tasks that take longer to master should also be considered to determine whether one specific learning type might be more useful at certain points in the learning process. The effect of different learning types on retention and generalization while acquiring complex skills is of interest as previous research has shown that learning from errors and through reinforcement can have different consequences on relearning and retention (Wood et al., 2024). Finally, it will be interesting to understand what actually gets reinforced while learning a complex task and to have graded feedback split by elements of the task.

## Conclusions

We found that the different learning types affect learning a complex skill like maze navigation in distinct ways. Specifically, the reinforcement feedback with an adapted criterion resulted in learning and transfer outcomes comparable to those of the group receiving error feedback. This suggests that reinforcement feedback might be equally useful to learn complex skill tasks. Additionally, our results show that repeated practice can reduce variability associated with transfer, although some form of feedback is necessary to significantly improve accuracy. Finally, we observed no changes in proprioception across any of the groups, this requires further research to determine whether and how proprioceptive adjustments might occur.

