## Supplementary figures and images for "The Impact of Different Learning Processes on Acquisition, Transfer, and Proprioception in Complex Motor Tasks"

### Supplementary Figure 1

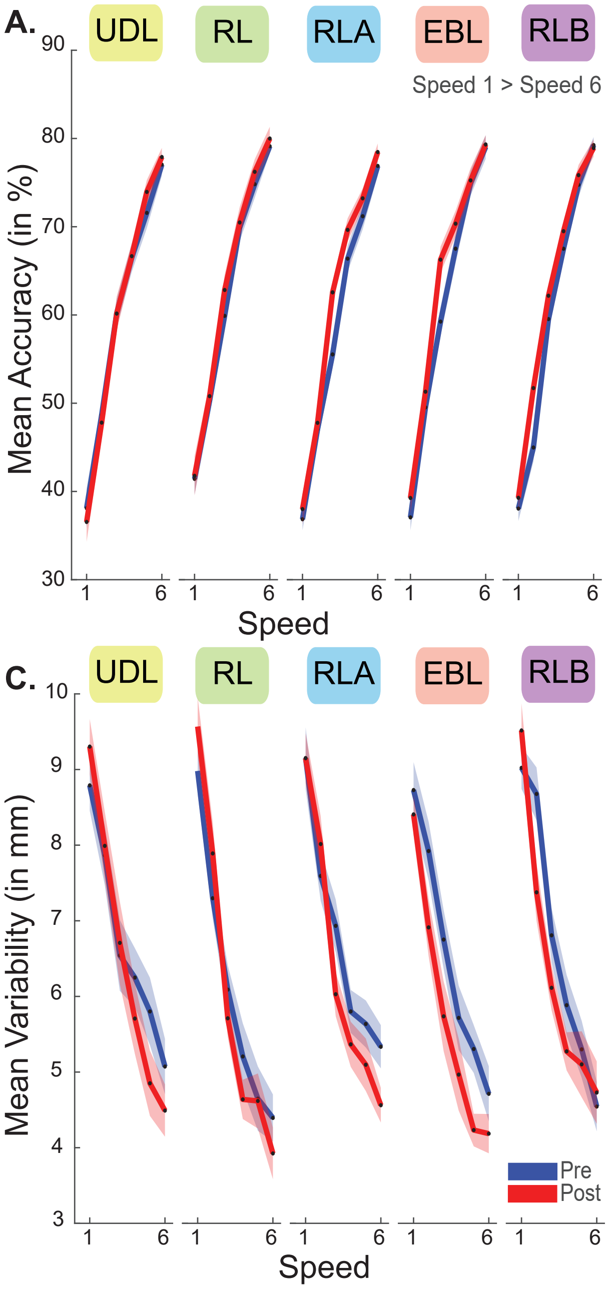

### Supplementary Figure 2

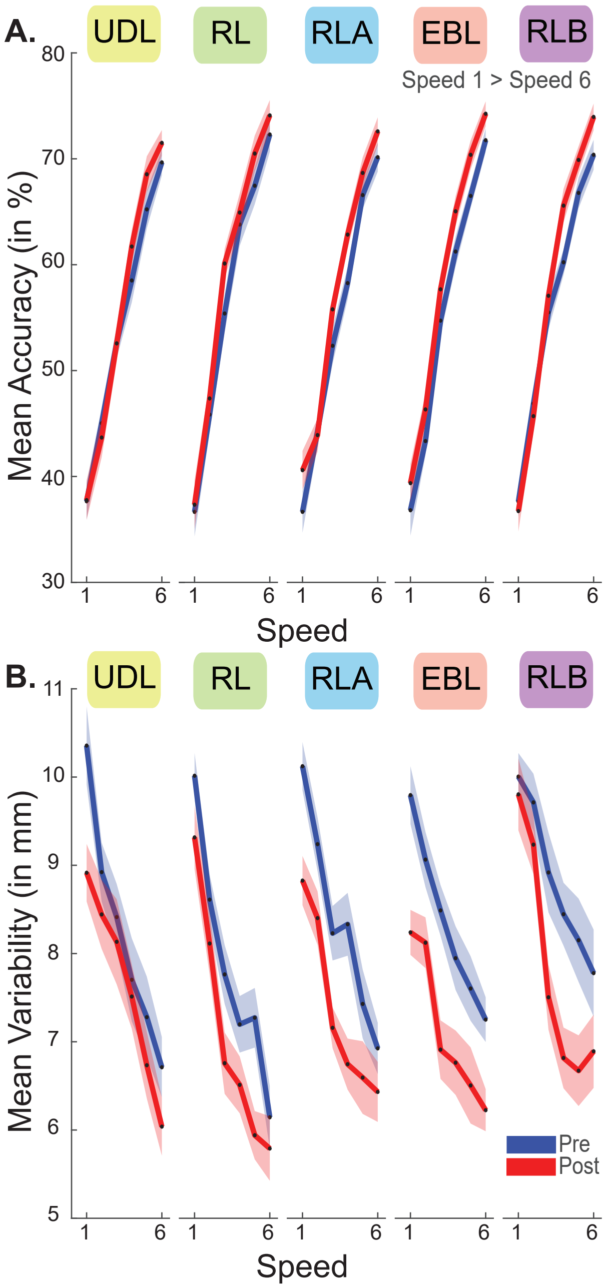
